# Effect of bedrest on the human gut and oral microbiome: implications for frailty

**DOI:** 10.64898/2025.12.21.695811

**Authors:** Monica Alvaro-Fuss, Vanessa DeClercq, Joanna M. Blodgett, Olga Theou, Morgan G.I. Langille, Robert G. Beiko

## Abstract

The physiological effects of spaceflight resemble those of ageing and prolonged inactivity, and ground-based microgravity analogs have emerged as promising models for studying frailty. The human microbiome is increasingly recognised for its role in age-associated decline, although precise mechanisms remain unclear. Here, we evaluate the gut and oral microbiomes of twenty-two participants, aged 55-65, who were enrolled in a head-down tilt bedrest (HDBR) study, the first Canadian HDBR study conducted in an older cohort. Participants were randomly assigned to an inactivity or multi-modality exercise intervention group for fourteen days of HDBR, followed by seven days of rehabilitation and additional follow-up appointments. Gut (n=343) and oral (n=344) taxonomic profiles were generated using V4/V5 16S rRNA gene sequencing from fecal and salivary samples collected throughout the study. Gut functional profiles were generated using metagenomic (n=86) and metabolomic (n=83) data. Frailty was measured using a 36-item frailty index. Inactivity-associated changes to the gut microbiome during HDBR included decreasing *α*-diversity, decreasing *Akkermansia* and *Lactobacillus*, and increasing *Bacteroides*. Exercise-associated changes included increasing gut *Roseburia*. Interestingly, oral microbiome *β*-diversity was more strongly associated with frailty than the gut. We conclude that inactivity-associated changes to the human microbiome could contribute to the early stages of frailty development, and that exercise can serve as an effective countermeasure against these effects. These results may inform strategies to preserve the health of both older adults facing prolonged periods of inactivity, as well as astronauts during longer space exploration missions.

**IMPORTANCE:** Healthy ageing carries significant demographic, economic, and public health implications globally, and both physical activity and human-associated microorganisms play important roles in its trajectory. Using microgravity simulation in older adults as a human frailty model, we show that short periods of inactivity could have significant effects on these microbial communities, particularly in the gut, which could further contribute to multi-system physiological decline and frailty pathogenesis. Participants who completed a daily, multi-modality exercise intervention did not exhibit these changes, which may help protect both older adults facing periods of inactivity and astronauts in the advent of longer space exploration missions. Interestingly, our results suggested that oral bacteria may be a more sensitive indicator of individual frailty levels than gut bacteria. Future research should ensure that functional aspects of human-associated microbial communities are thoroughly investigated, as these may offer deeper insight into their contributions to age-related decline.

## INTRODUCTION

The human gastrointestinal tract harbours a vast network of microbial life [1]. These microbial communities, particularly gut-associated ones, are increasingly recognized as integral to host physiology, influencing multiple organs and systems through various host–microbiome axes [2, 3, 4, 5, 6, 7]. Community structure is shaped by diverse physicochemical and nutritional gradients [8, 9], determined by environmental, lifestyle, and genetic factors [10, 11], and shifts in microbiome composition during ageing are thought to contribute to physiological decline in older adults [12, 13, 14]. In the gut, frailty and non-communicable diseases are associated with reduced microbial *α*-diversity, lower metabolic capacity, loss of beneficial taxa, and increased abundance of opportunistic pathogens, which are also linked to states of chronic low-grade inflammation and immunosenescence [15, 16, 17, 18, 19]. Moreover, the role of the oral microbiome in local and systemic health, including potential links to non-communicable diseases, have also gained increasing attention in the literature [14, 20, 21]. However, the full range of mechanisms through which microbial communities shape human ageing remains to be elucidated.

Frailty refers to a state of increased vulnerability to adverse health outcomes resulting from age-related decline across multiple physiological systems [22]. Clinical measures, such as frailty indexes, are used to measure this vulnerability, helping to differentiate biological from chronological ageing [22, 23]. With rising life expectancy and globally ageing populations, promoting healthy ageing carries significant demographic, economic, and public-health implications [24, 25]. In recent decades, advancements in space exploration research have uncovered intriguing parallels between the effects of microgravity and those of biological ageing [26, 27], suggesting that microgravity analogs could be leveraged for studying frailty in humans. A review conducted by our group in 2019 showed that 6^◦^ head-down tilt bedrest (HDBR), a common ground-based method used to reduce gravitational stimuli, increases frailty-related biomarkers over short periods of time, with notable effects on the cardiovascular and musculoskeletal systems [28]. Further, the effects of HDBR resemble those of prolonged inactivity [26, 27], which is thought to contribute to at least 35 age-associated non-communicable diseases and can accelerate biological ageing [29]. As a result, exercise has been established as a primary countermeasure to mitigate physical deconditioning since early human spaceflight [30].

Physical activity is a cornerstone of healthy ageing [31, 32, 33]. It supports the function of many biological systems, slows down ageing markers, and reduces the onset and progression of frailty and non-communicable diseases [34, 35, 36]. Growing evidence suggests that physical activity also affects the composition of human-associated microbial communities, particularly in the gut. Exercise is shown to promote gut microbiome *α*-diversity, abundance of health-associated taxa, and production of short-chain fatty acids (SCFAs), a key metabolic output of gut microbiota with important local and systemic effects [37, 38, 39, 40]. Murine models further suggest that low-to-moderate intensity exercise also plays a role in preserving intestinal barrier function and preventing permeability [41, 42, 43], which is essential for regulating host–microbiome interactions, including immune responses [44]. Although less is known about the relationship between the oral microbiome and exercise, emerging evidence indicates that physical activity can indeed influence its diversity and composition, and plays important roles in nitric oxide metabolism with implications for cardiovascular and musculoskeletal function [45, 46, 47].

In addition, the duration, type, and frequency of physical activity, along with factors such as diet, can influence the extent and nature of responses in both the host and host-associated microbial communities [48, 49, 50].

The present study, which was supported by the Canadian Institutes of Health Research, the Canadian Frailty Network, and the Canadian Space Agency, was the first Canadian HDBR study conducted in mid-older aged adults [51]. This interdisciplinary initiative, involving eight research teams across Canada, was a two-arm randomized controlled trial that investigated the physiological effects of fourteen days of HDBR with or without a multi-modality exercise countermeasure [30]. Here, we evaluated the gut and oral microbiome of study participants to explore links between inactivity, exercise, frailty, and human-associated microbial communities. We analyzed a total of 343 fecal and 344 salivary samples using 16S rRNA gene sequencing to generate taxonomic profiles, and further generated functional profiles of the gut microbiome using 86 metagenomic and 83 metabolomic samples. We hypothesized that inactivity would induce frailty-related disruptions in microbial communities, particularly in the gut, while exercise would have protective effects on microbiome structure. Indeed, we show that inactivity-associated changes to gut microbiome structure during HDBR could contribute to increased frailty. Our results may inform strategies that promote healthy ageing in older adults, as well as contribute to the development of microbiome-informed countermeasures to safeguard astronaut health in anticipation of longer-duration space missions.

## MATERIALS AND METHODS

### Study design

An in-depth overview of the study protocol, including participant recruitment criteria and exercise countermeasure details, is provided in [51]. Briefly, twenty-two healthy participants were enrolled at the McConnell Centre for Innovative Medicine, Research Institute of the McGill University Health Centre in Montreal, QB between July and December 2021. Participants were randomized into the control (inactivity) or exercise groups, in four separate cohorts of five or six individuals. They were free to circulate for the initial five-day adaptation period prior to the fourteen-day HDBR phase. During the HDBR phase, the control group carried out stretch and motion physiotherapy exercises, while the exercise group completed a structured, multi-modality daily exercise program consisting of high-intensity interval training, low-intensity aerobic training, and both lower and upper limb training. All participants could freely toss, turn and stretch in bed and were kept on a weight-maintaining diet. All activities, including eating, bathing, toileting and exercise, were carried out in the HDBR position. Day and night cycles were standardised such that participants were awake and sleeping at the same time throughout bed rest (awake from 7:00am to approximately 11:00pm to allow for a 16-hour awake cycle). Following HDBR, the participants underwent a seven-day recovery period where they were rehabilitated and allowed to ambulate under supervision. The exercise intervention was not continued during the recovery period. Participants returned to the facility four weeks and four months after the in-patient period for follow-up appointments.

### Frailty measurement collection

Frailty was measured using a symptom-based 36-item frailty index (FI-36) at baseline, days 1, 3, 7, 12 and 14 of HDBR, at days 2 and 6 of recovery, and both follow-up appointments. Ascertainment of the FI-36 followed established protocols [52] and has been published elsewhere [Blodgett *et al., under review*]. Briefly, the FI-36 was derived as the number of deficits present divided by the number of deficits considered (e.g. 18 deficits out of 36 considered = FI score of 0.5). The FI-36 consisted of 33 symptom-based deficits and three self-reported health questions. The 33 symptoms were coded as binary 0-1 deficits, while the three self-reported measures (vision, hearing, general health) were coded as 0, 0.25, 0.5, 0.75 or 1. Therefore, the maximal range of FI-36 scores are between 0 and 1, with higher scores indicating greater frailty levels.

### Fecal and salivary DNA sample collection and extraction

Stool and saliva samples were collected from participants once during the adaptation period, daily during bedrest, at days 3 and 6 of recovery, and at both follow-up appointments. Fecal samples were collected from sterile bedpans using OMNIgene GUT collection and stabilization kits (OMR-200) as soon as possible after the bowel movement.

Saliva samples were collected using OMNIgene SALIVA DNA and RNA device (OMR-610). All samples were frozen and stored at -20^◦^C, then sent to the Integrated Microbiome Resource at Dalhousie University and stored at -80^◦^C until processing. DNA was extracted from all fecal and salivary samples using the QIAGEN PowerFecal DNA Kit (cat: 51804).

### 16S rRNA gene amplicon sequencing and bioinformatics

The V4-V5 region of the 16S rRNA gene was PCR-amplified with universal 515FB (GTGYCAGCMGCCGCGGTAA)/926R (CCGYCAATTYMTTTRAGTTT) primers and sequenced on an Illumina MiSeq using 300+300 bp paired-end V3 chemistry. Amplicon data were analyzed as per Microbiome Helper Amplicon SOP v2 [53] using QIIME2 version 2022.8 [54]. Primer sequences were removed using the cutadapt plugin [55] and untrimmed reads were discarded. Trimmed reads were denoised using the DADA2 plugin [56]. Amplicon sequence variants (ASVs) with an abundance lower than 0.1% of the mean sample depth across all samples were discarded due to the risk of sequencing instrument bleed-through [57]. Taxonomy was assigned using the Naïve Bayes classifier implemented in the Scikit-learn library plugin [58, 59] against the SILVA-138 database [60]. ASVs mapping to mitochondrial or chloroplast sequences, or those that remained unclassified at the phylum level, were removed from the analysis. Phylogenetic distances between ASVs were determined via fragment insertion using SEPP [61]. Sample *α*- and *β*-diversity were calculated at the ASV level. To calculate sample *α*-diversity, we first generated 100 random subsamples of gut and oral feature tables, rarefied to 2,500 reads. We then calculated mean observed richness, mean Shannon entropy [62], and mean Faith’s phylogenetic diversity [63]. *β*-diversity was calculated on unrarefied ASV tables using phylogenetic robust Aitchison PCA implemented in the gemelli toolbox [64].

### Fecal metagenomic sequencing and bioinformatics

Metagenomic sequencing was performed on stool samples collected on days 1, 7 and 14 of bedrest, as well as day 6 of recovery, matching samples to those collected for metabolomic analysis. Metagenomic DNA was processed using the Illumina Nextera Flex kit and sequenced on an Illumina NextSeq 2000 using 150+150 bp paired-end "high output" chemistry. Metagenomic data were analyzed as per Microbiome Helper Metagenomics SOP v2 and v3 [53]. Raw paired sequencing reads were processed using Kneaddata as implemented in BioBakery3 [65] at default parameters and concatenated into a single file. Metabolic pathway abundance was determined using HUMAnN3 [65] at default parameters, and unstratified MetaCyc pathway abundance data was used for downstream analysis.

### Fecal metabolomic sample collection and analysis

Stool samples for metabolomics were collected from sterile bedpans using OMNImet GUT collection and stabilization kits (ME-200) as soon as possible after the bowel movement on days 1, 7 and 14 of bedrest, and day 6 of recovery. All samples were frozen and stored at -20^◦^C, then sent to the Integrated Microbiome Resource at Dalhousie University and stored at -80^◦^C until processing. Untargeted metabolomics were carried out by The Metabolomics Innovation Centre (TMIC) in Edmonton, AB as per their Microbiome Metabolome Assay. Data were normalized as a ratio of total useful signal by the TMIC as part of the analysis. Only metabolites categorized as microbiota-related (i.e. not host-microbiota-related) and belonging to quality Tier 1 (positive identification) were used for downsteam analysis.

### Statistical analysis

Statistical analysis was performed in R (version 4.3.2) using tidyverse [66]. Group differences at baseline in body mass index (BMI) and frailty were calculated using Kruskal-Wallis tests. Multivariate *β*-diversity associations were analyzed using PERMANOVA tests at 999 permutations as per vegan::adonis() [67]. Longitudinal *α*-diversity and differential abundance analyses were performed using linear mixed-effects models implemented in lme4::lmer() [68]. The effect of day of HDBR on *α*-diversity was analysed using the model diversity ∼ cohort + sex + day + (1 + day | participant_id). Sex interaction effects on *α*-diversity during HDBR were analyed using the model diversity ∼ cohort + sex ∗ day + (1 + day | participant_id). Prior to calculating differential abundance, all genus and pathway count data were transformed using the centered log-ratio (CLR) [69] with phyloseq [70] and microbiome::transform [71]. After CLR transformation, data were filtered to remove features with ≤10% prevalence across samples. The effect of day of HDBR on feature abundance was analyzed using the model abundance ∼ cohort + sex + day + (1 | participant_id). P-values derived from mixed-effect models were determined using Satterthwaite’s method [72, 73] for calculating degrees of freedom with lmerTest::summary() [74]. Differential abundance analysis p-values were adjusted as per Benjamini-Hochberg’s false discovery rate correction using rstatix::adjust_pvalue().

## RESULTS

### Participant overview and microbial community profiling

Data from all twenty-two participants that completed the HDBR phase were retained for analysis. Two participants (PT26 and PT27) withdrew during the recovery period due to atrial fibrillation. Mean participant age was 59.2 ± 2.94 years. BMI was calculated from height and weight measurements, and frailty was measured throughout the study as FI-36. There were significant differences in mean body mass index (BMI) between combined sex and exercise groups (*H*(3) = 0.418; *p* = 0.015). BMI appeared to be higher in males than females among both groups. There was no significant difference in baseline frailty between groups (*p* = 0.17) (Table 1). Taxonomic profiles of the gut and oral microbiomes of study participants were generated using 16S rRNA gene sequencing of the V4-V5 region on 343 fecal and 344 salivary samples collected throughout the study. Samples were only collected on days when a bowel movement occurred (Fig. S1, Fig. S2). Gut and oral relative abundance at the family level are shown in Fig. 1a.

**FIG 1.**
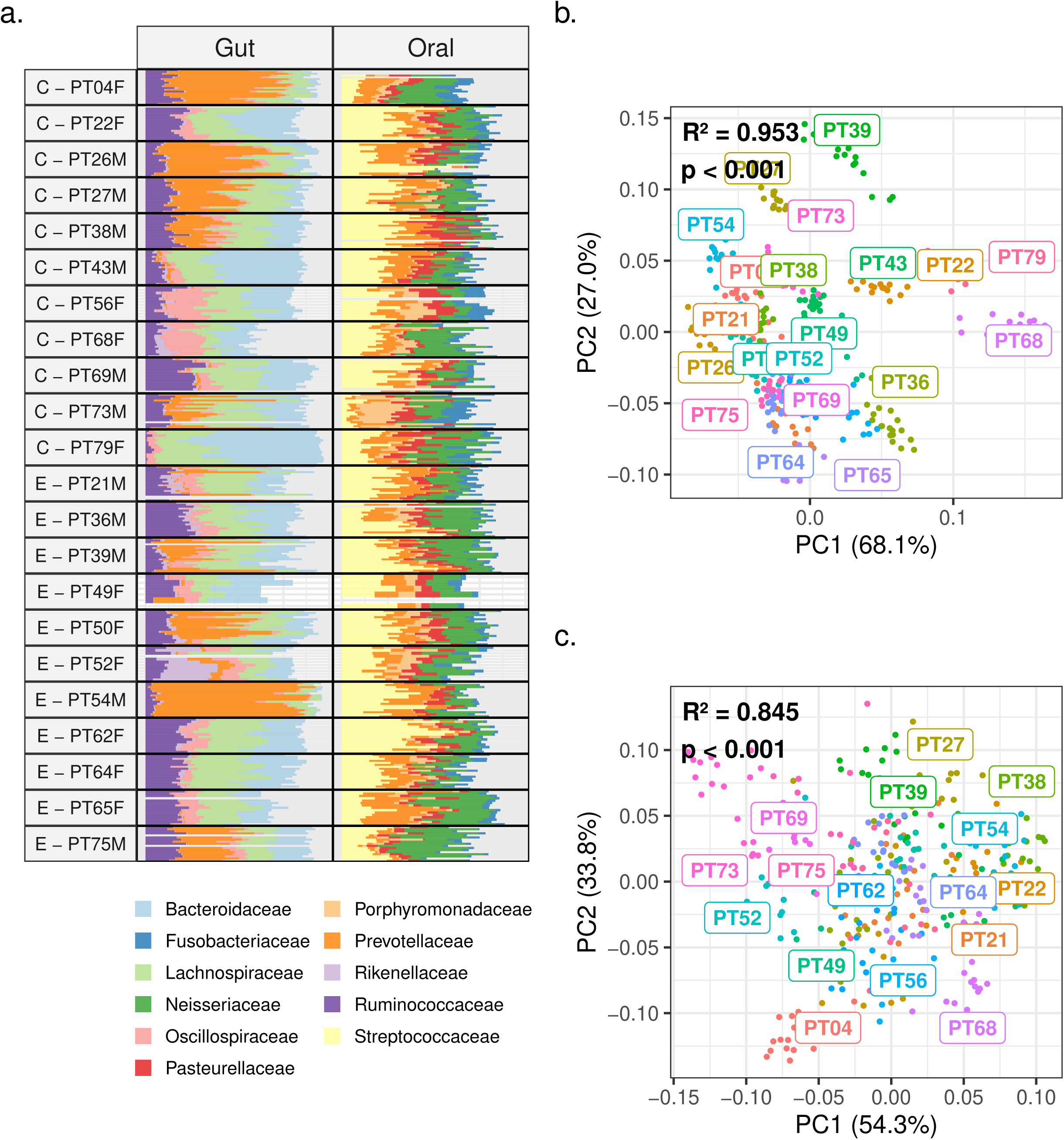
a) Taxonomic microbiome profiles of study participants (C: control; E: exercise; F: female; M: male). ASV relative abundances were aggregated at the family level. b–c) Ordination plots of b) gut and c) oral microbiome *β*-diversity, calculated using phylogenetic RPCA at the ASV level. Multivariate associations were assessed using PERMANOVA. Different colours represent each participant.

**TABLE 1.** Characteristics of CAIS participants grouped by intervention and sex.

|  | Control (F) | Control (M) | Exercise (F) | Exercise (M) |
| --- | --- | --- | --- | --- |
| *n* | 5 | 6 | 6 | 5 |
| Age (years) | 56.6 ± 1.14 | 60.8 ± 3.66 | 60.7 ± 2.34 | 58.0 ± 1.73 |
| Height (cm) | 161 ± 4.51 | 171 ± 12.7 | 161 ± 3.61 | 175 ± 4.45 |
| Weight (kg) | 58.6 ± 6.95 | 76.0 ± 14.7 | 63.0 ± 8.93 | 84.9 ± 5.42 |
| BMI ($kg/m^2$) | 22.5 ± 2.34 | 25.8 ± 2.16 | 24.4 ± 3.21 | 27.8 ± 1.03 |
| Baseline FI-36 | 0.038 ± 0.027 | 0.031 ± 0.019 | 0.028 ± 0.031 | 0.0083 ± 0.012 |

<sup>a</sup>All values are presented as mean ± SD. F: female; M: male.

### Associations between microbial *β*-diversity and study variables

To investigate associations between participant microbiome profiles and various study variables, we performed PERMANOVA tests against community *β*-diversity. *β*-diversity was calculated as phylogenetic RPCA at the ASV level. *β*-diversity of both gut and oral samples was strongly associated with participant ID (Fig. 1b, Fig. 1c). Participant sex, cohort and exercise group were also significantly associated with sample *β*-diversity, although effect sizes were small (Fig 2). We further investigated associations between gut and oral microbiome *β*-diversity and FI-36 measurements (gut: n = 152; oral: n = 154).

**FIG 2.**
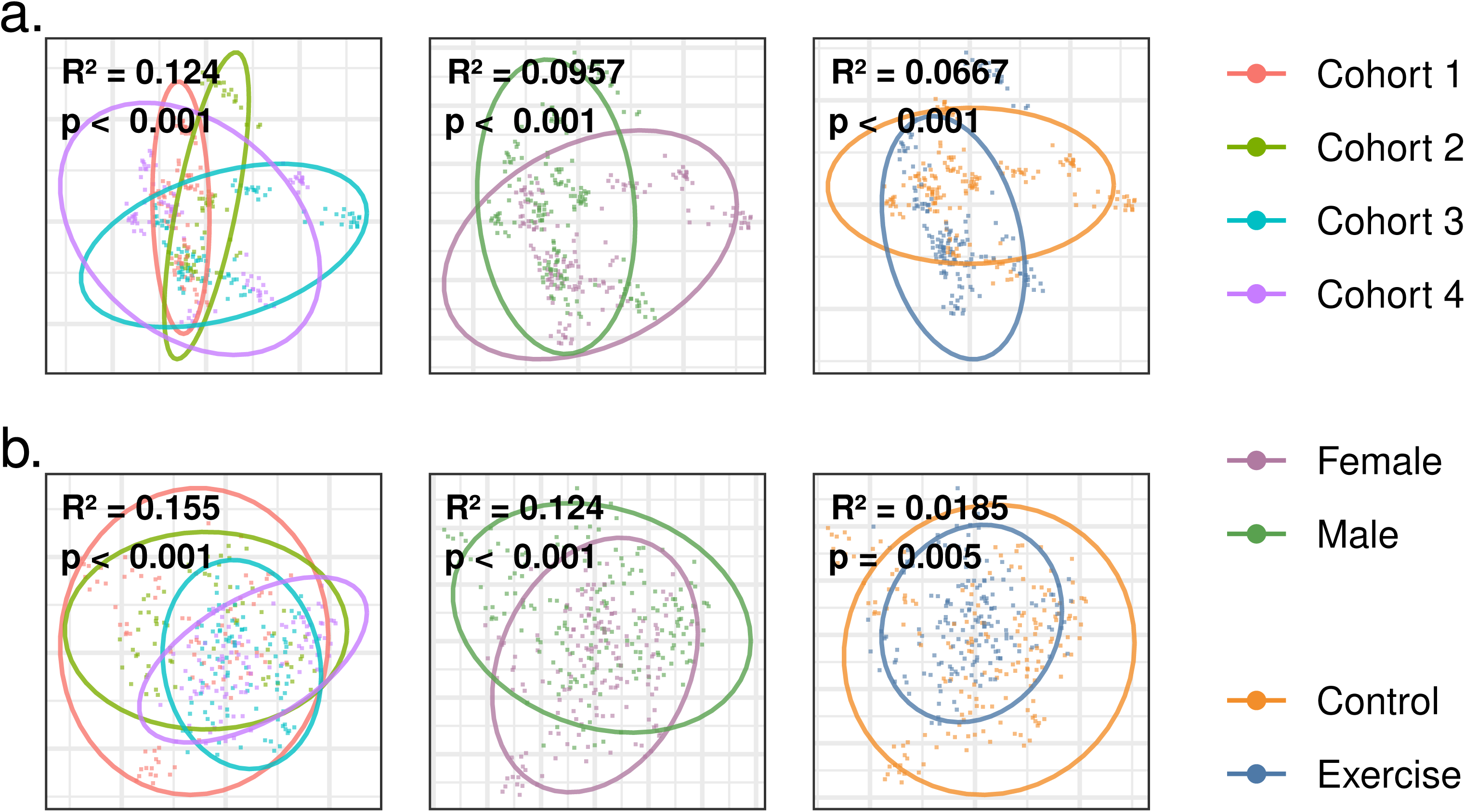
Ordination plots of a) gut and b) oral microbiome *β*-diversity, calculated using phylogenetic RPCA at the ASV level. Multivariate associations were assessed using PERMANOVA.

Oral microbiome *β*-diversity (*R*^2^ = 0.0649, *p* = 0.001) showed a stronger association with frailty than the gut microbiome (*R*^2^ = 0.00855, *p* = 0.28) in the uncorrected model. After adjusting for participant ID, both associations were non-significant (p > 0.05), though the oral microbiome remained comparatively stronger (Fig. 3). Removing a single high-frailty outlier (FI > 0.6) strengthened both associations, with oral *β*-diversity regaining statistical significance (gut: *R*^2^ = 0.00038, *p* = 0.43; oral: *R*^2^ = 0.00442, *p* = 0.035).

**FIG 3.**
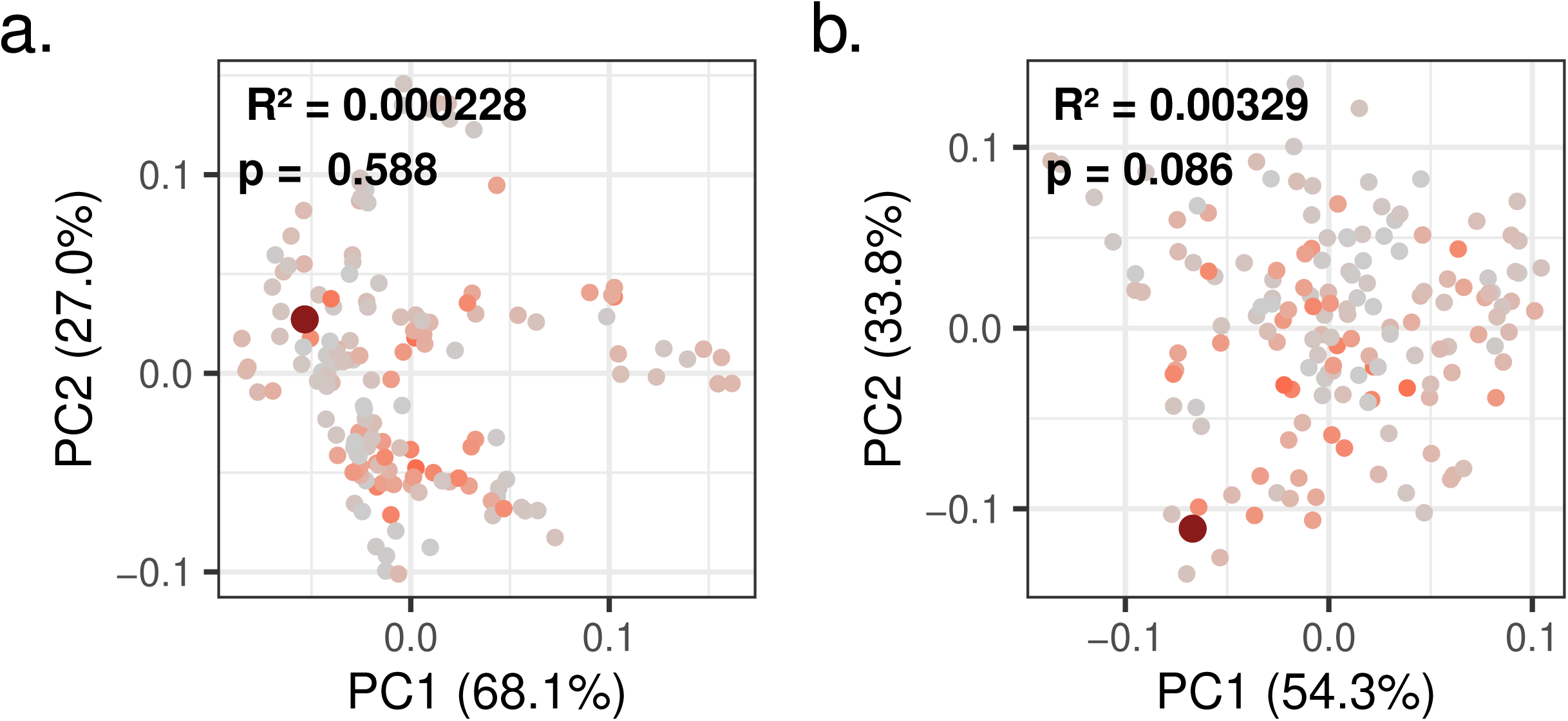
Ordination plots of a) gut and b) oral microbiome *β*-diversity, calculated using phylogenetic RPCA at the ASV level. High frailty outlier (FI-36 > 0.6) is highlighted in dark red. Multivariate association with frailty (FI-36) was assessed using PERMANOVA. *R*^2^ and p values are adjusted for participant ID.

### Decreasing gut microbiota alpha diversity is associated with inactivity during HDBR

Gut and oral microbiome *α*-diversity was measured using observed richness, Shannon entropy, and phylogenetic diversity after performing repeated rarefaction. To evaluate longitudinal changes in *α*-diversity during HDBR associated with inactivity and exercise, we applied linear mixed models to exercise-stratified data. All gut *α*-diversity metrics decreased during HDBR in the control group, but did not change in the exercise group (Fig 3, Table S2). We did not identify any interaction effects with sex (Table S3). Due to limited timepoints, we did not carry out statistical analysis of *α*-diversity during the recovery phase, although visual representation of the data suggested that gut *α*-diversity of the control group increased again after HDBR (Fig 3a).

### Differential abundance of taxa, pathways and metabolites during bedrest

To further investigate the effects of inactivity and exercise on microbiome composition, we carried out longitudinal differential abundance analysis of microbial genera during HDBR using linear mixed models. The control group exhibited more changes in gut and oral microbial genera than the exercise group. The control group exhibited decreases in gut *Akkermansia*, Bacilli RF39, *Lactobacillus*, Oscillospiraceae UCG-003, *Rumicococcus gauvreauii group*, Clostridia UCG-014 and *Subdoligranulum*, as well as increases in an unclassified Barnesiellaceae, *Howardella*, *Alloprevotella*, *Prevotella* NK3B31 group, *Paraprevotella*, *Bacteroides* and Lachnospiraceae *UCG-003*. Conversely, the exercise group exhibited decreases in gut *Terrisporobacter* and *Turicibacter*, as well as increases in gut *Roseburia* and *Clostridium* sensu stricto 1 (Figure 4a). The oral microbiome of the control group exhibited increases in *Rothia* and decreases in *Lautropia* and *Abiotrophia*. The oral microbiome of the exercise group exhibited increases in an unclassified Comamonadaceae and *Pseudopropionibacterium* (Fig 4b). Abundance of some, but not all taxa, exhibited recovery trajectories after HDBR.

**FIG 4.**
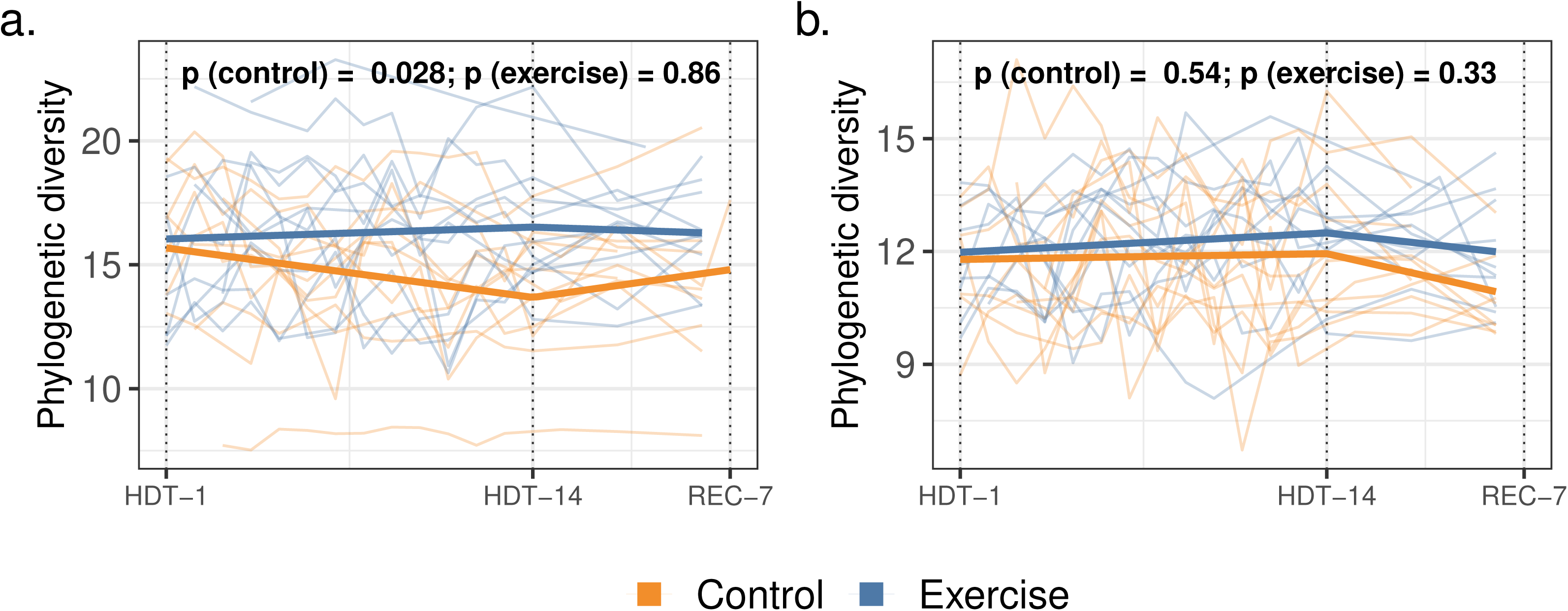
a) Gut and b) oral microbiome *α*-diversity trajectories during HDBR and recovery. Background lines represent individual participants. Bold lines indicate group-level trends. Dotted lines indicate beginning of bedrest (HDT-1), end of bedrest (HDT-14) and end of recovery (REC). Change in *α*-diversity during HDBR was analysed using mixed-effect models.

**FIG 5.**
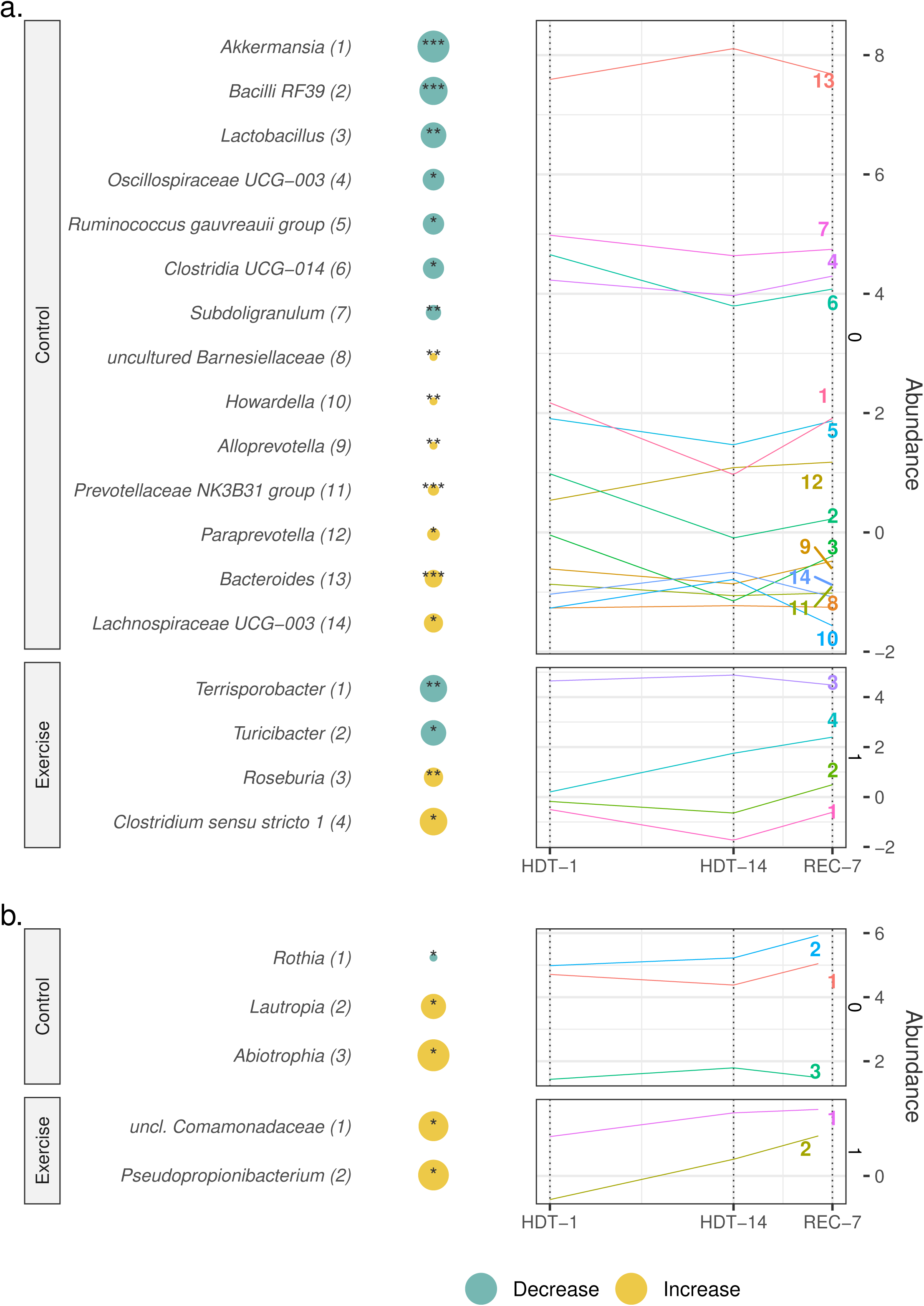
Longitudinal differential abundance analysis of a) gut and b) oral microbial genera. Circle size indicates the magnitude of the change in abundance during HDBR. Adjusted p-values are represented as (p < 0.001: ***; p < 0.01: **; p < 0.05: *). Right hand panels show the grouped CLR abundance of differentially abundant taxa across bedrest and recovery. Dotted lines indicate beginning of bedrest (HDT-1), end of bedrest (HDT-14) and end of recovery (REC). Individual features are represented by their corresponding number. Change in feature abundance during HDBR was analysed using mixed-effect models.

To further evaluate functional changes in the gut microbiome during HDBR, we carried out metagenomic and metabolomic analysis of select fecal samples (Fig S5). Metagenomic sequencing data were used to infer the abundance of MetaCyc metabolic pathways. Longitudinal differential abundance testing of metabolic pathways and metabolites in both the control and exercise groups was performed using the same linear mixed model framework. After applying false discovery rate correction, no pathways or metabolites remained as statistically significant (adjusted p < 0.05). Differential abundance results based on raw p-values are shown in Fig S4 and S5. Briefly, pathways possibly decreasing in the control group included heterolactic fermentation, tetrahydrofolate biosynthesis, D-ravidosamine biosynthesis, palmitate and myristate biosynthesis, and cysteine biosynthesis and interconversion. Possibly increasing pathways included both citrulline and 5-oxoproline metabolism, as well as both pyridoxal-5-phosphate and citrulline biosynthesis. In the exercise group, possibly increasing pathways included nitrate and sulfate assimilation pathways, and D-ravidosamine biosynthesis. Possibly decreasing metabolites in the control group included phenylpyruvate and 3-methyladipic acid. Possibly increasing metabolites in the control group included hypotaurine and several trytophan catabolites, such as indole-3-propionic and indole-3-acetic acid. In the exercise group, possibly decreasing metabolites included cholic acid, glycocholic acid, and several secondary bile acids. Possibly increasing metabolites in the exercise group included phenylacetylglutamine and syringic acid.

## DISCUSSION

Microgravity analogs hold potential as human models for studying frailty, given similarities with the physiological effects to ageing and prolonged inactivity [26, 27, 28]. The present study examined the effects of 14 days of HDBR with or without exercise on healthy mid-older adults, and was the first Canadian study of its kind [51]. Hajj-Boutros *et. al.* reported that body weight fell in both groups, but exercise preserved lean mass and aerobic fitness [75]. Blodgett *et. al.* recorded that participant frailty increased during bedrest, decreased during recovery, and did not return to baseline levels after four months, and that those in the exercise group exhibited better recovery trajectories *(unpublished)*. Additional results can be found in [76, 77, 78]. Here, we assessed participants’ gut and oral microbiomes to provide insight into the frailty-microbiome relationship. Differences in community profiles were mostly driven by subject-specific signatures, supporting the idea of highly individualised microbiomes [10, 11]. Participant cohort, sex and exercise group were also associated with community composition, although effect sizes were markedly lower. Furthermore, longitudinal analysis revealed several changes to gut and oral microbiome structure associated with both inactivity and exercise during HDBR, with notable links to the existing frailty-microbiome literature.

Gut microbiome *α*-diversity plays a key role in maintaining cooperative cross-feeding networks [79, 80]. Decreases in *α*-diversity impact the availability of key metabolic intermediates [79], reduce a community’s resilience to selective stressors [81, 82, 83], and are commonly associated with frailty and non-communicable diseases in older adults [12, 13, 84, 85]. Conversely, higher gut microbiome *α*-diversity is often observed in athletes and physically active men and women, correlating with improved cardiorespiratory and metabolic health [86, 87, 88]. Consistent with these findings, our results suggest that inactivity during HDBR led to reductions in gut microbiome *α*-diversity independently of sex, whereas this was not observed with the daily exercise intervention. Several mechanisms have been proposed to explain how physical activity may increase gut microbiome *α*-diversity, such as decreased stool transit time, tissue hypoxia, improved immune function, and enhanced barrier integrity [48, 89, 90].

Additionally, elevated mitochondrial reactive oxygen species production, which is strongly related to frailty, immunosenescence, and biological ageing [91, 92], was further associated with reduced gut microbiome *α*-diversity in a 2019 study [93].

Physical activity promotes gut microbiome-derived SCFA production, particularly butyrate [94, 95]. Butyrate is essential for colonocyte metabolism, supports intestinal barrier integrity, and helps suppress inflammatory responses [96, 97, 98]. Reduced butyrate and other SCFAs have been linked to frailty and numerous non-communicable diseases across physiological systems [12, 13, 15, 16, 17]. *Subdoligranulum*, which decreased in the control group, and *Roseburia*, which increased in the exercise group, are considered butyrogenic genera [99, 100, 101], with *Roseburia* abundance often reported to rise with physical activity [101, 89, 102]. Furthermore, butyrate production is largely regulated by non-butyrogenic taxa, particularly through cross-feeding with acetate [79, 103, 104]. The genus Bacilli RF39, which decreased in the control group, has been identified as a primarily acetogenic taxon and may contribute to this regulation [105]. These results suggest that inactivity impacts butyrate production both directly and indirectly, with possible local and systemic implications. While several members of the Bacteroidales order, which also include many SCFA-producing species, increased in the control group, certain taxa such as *Bacteroides* and *Paraprevotella* have been associated with unhealthy ageing and extraintestinal infections [106, 107, 108, 109].

Changes in gut microbiome composition can also impact intestinal barrier function [8]; delimitation of such organism-intrinsic barriers is considered a hallmark of health [110]. The genus *Akkermansia*, which decreased in the control group, plays an important role in barrier health through mucin degradation [111, 112] and positively correlates with *α*-diversity and healthier metabolic status [112, 113]. The genus *Lactobacillus*, which also decreased with inactivity, is thought to have protective immunomodulatory effects [114], and contributes to butyrate biosynthesis through lactic acid production [115, 79].

Increased intestinal permeability contributes to frailty and age-associated non-communicable diseases [12, 13], and can also affect microbiome structure. Diffusion of host oxygen into the anaerobic luminal environment can interfere with butyrate synthesis and promote the growth of opportunistic pathogens [96]. In fact, some frameworks have suggested that increased luminal oxygenation is a main driver disease-associated gut microbiome shifts [116, 117]. Gut *Akkermansia* and *Lactobacillus* have been often reported to increase with exercise interventions in both human and murine models [118, 119, 120, 121, 122, 123], and also been found to decrease in other inactivity-based microgravity analog studies [124, 125] Functional analysis of the gut microbiome did not yield statistically significant results, likely due to the smaller sample size compared with the 16S rRNA gene dataset. Further, changes in pathway abundance do not necessarily reflect changes in pathway activity.

Nonetheless, results based on raw p-values may align with and strengthen our findings. Heterolactic fermentation, the pathway most reduced in the control group, contributes to butyrate synthesis through lactate-mediated cross-feeding [79, 126, 127]. Loss of gut *α*-diversity has been linked to reduced tetrahydrofolate biosynthesis [79], which may also decline with inactivity and is strongly associated with ageing [128]. Signs of antioxidant activity, such as increased 5-oxoproline metabolism and higher hypotaurine abundance, were observed with inactivity, potentially reflecting elevated luminal oxygenation [129, 130, 131]. Although SCFA levels and biosynthetic pathways were not differentially abundant, antioxidant presence may sustain butyrate production under aerobic conditions [132]. Moreover, longer-duration HDBR studies such as the AGBRESA study have indeed reported changes to SCFA profiles [133]. Inactivity was also associated with reduced levels of the tryptophan catabolites indole-3-propionic and indole-3-acetic acid, which exert intestinal and systemic anti-inflammatory and antioxidant effects and respond to barrier damage [134, 135, 136, 137]. However, sustained activation of their receptor, the aryl hydrocarbon receptor, is linked to immunosenescence and disease pathology [138].

Less is known about the oral microbiome, but emerging evidence suggests these communities also play important roles in systemic health and frailty [20, 21, 14]. One study reported that fifteen days of HDBR was associated with increased microbial richness and shifts in certain genera [139], but we did not observe similar diversity changes or alterations in the same taxa. However, we found that oral microbiome *β*-diversity was more strongly associated with frailty than gut microbiome *β*-diversity, despite the latter being more commonly linked to the underlying mechanisms of frailty. While the effect size was low, this suggests that the oral microbiome may serve as a compelling indicator of individual frailty levels, particularly given its easier and less invasive sampling compared to the gut. The oral microbiome has also been proposed as a diagnostic and prognostic biomarker for oral and colorectal cancers, with attention often drawn to specific oral pathogens such as *P. gingivalis* and *F. nucleatum* [140, 141, 142, 143]. Oral health and hygiene likely play a significant role in shaping the oral microbiome–frailty relationship, as individuals with lower functional capacity may engage less in routine practices such as toothbrushing [144]. Unfortunately, oral health or hygiene data were not collected before or during the study.

This study had several limitations. The strict inclusion and exclusion criteria resulted in a cohort of healthy volunteers, which may limit the generalizability of our findings to more diverse populations. Detailed health records, including pre-study exercise and dietary habits, were unavailable, preventing a more comprehensive assessment of lifestyle factors influencing microbiome composition; however, the two-arm randomized control trial study design may have helped address these challenges. While diet and exercise were standardized during the study, variations may still have contributed to some of the observed microbiome shifts. Future studies should include detailed dietary and activity tracking to better contextualize microbiome changes in response to prolonged inactivity and exercise interventions. Limited sample size and availability also reduced statistical power, particularly for metagenomic, metabolomic, and recovery-phase analyses. Linear mixed models helped manage missing and irregularly-spaced samples during HDBR. Consistent daily sampling in future studies would enhance statistical power and provide finer longitudinal resolution.

In conclusion, physical inactivity during bedrest in older adults induces shifts in microbiome structure that may represent early steps in frailty development, potentially triggering a self-reinforcing cycle of microbiome disruption and host decline centered on impaired intestinal barrier function. Daily multi-modality exercise mitigated these effects, reinforcing the protective role of physical activity. Notably, oral microbiome structure was more strongly associated with frailty than the gut microbiome; while the underlying reasons remain unclear, the oral microbiome is more accessible than the gut, and may warrant further investigation. These findings may inform strategies to support the health of older adults during inactivity and astronauts during spaceflight. Future studies should consider including larger cohorts, extended monitoring, and daily sampling to better capture temporal microbiome dynamics.

## ACKNOWLEDGMENTS

Financial support was provided by the Nova Scotia Graduate Scholarship.

## DATA AVAILABILITY STATEMENT

Data may be made available by the corresponding author upon reasonable request, as they may only be shared for the use under which they were ethically approved. Analysis code can be found at https://github.com/malvarofuss/CSA-inactivity-microbiome.

## CLINICAL TRIALS

The study was registered as a clinical trial (NCT04964999: Microgravity Research Analogue (MRA): Understanding the Health Impact of Inactivity for the Benefit of Older Adults and Astronauts Initiative) in the US National Clinical Trial Registry.

## ETHICS APPROVAL

Ethical approval for this study was obtained from the McGill Hospital University Centre Research Ethics Board (REB; MP-37-2021-7170), with project-specific approval granted by Dalhousie University REB (2021-5478). Participants signed a written informed consent and agreed to be available at the McGill Hospital University Centre for the entire 26-day in-patient period. The research was conducted in compliance with the guidelines and regulations of the above agencies and the declaration of Helsinki.

## Supplemental material

Supplemental tables and figures can be found in supplemental.pdf.

